# Biodegradation Potential of Neptunomonas naphthovoran - NAG-2N-126 for Crude Oil Pollution Mitigation: Experimental and Modelling Insights

**DOI:** 10.1101/2024.05.31.596902

**Authors:** Turan Mutallimov, Kejian Wu, Hai Deng, Yusif Ibrahimov

## Abstract

Crude oil contamination poses a significant threat to the environment, demanding effective bioremediation strategies. While microbial biodegradation holds promise, there is a notable absence of exploration, particularly regarding the biodegradation potential of Neptunomonas naphthovoran strain NAG-2N-126. This study addresses this gap by investigating the bioremediation potential of NAG-2N-126 through experimental analyses and mathematical modelling. Our findings reveal the strain’s rapid growth on crude oil, with the microbial biomass predicted to undergo a 3.5-fold increase over the experimental period. Additionally, the strain exhibited high efficiency in degrading aromatic compounds, achieving an impressive 41% degradation rate within the initial 20-day period. Oxygen availability emerged as a critical factor influencing microbial growth and biodegradation processes, with simulations indicating a 20% decrease in degradation rates under oxygen-deprived conditions. Mathematical modelling provided insights into the complex dynamics involved, contributing to the understanding of microbial hydrocarbon degradation. However, further research is warranted to optimize growth conditions and scale up findings for practical applications in environmental cleanup efforts. By delving into this underexplored area, this study highlights the importance of NAG-2N-126 in crude oil bioremediation and underscores the need for further research to optimize its application in environmental cleanup efforts.

**IMPORTANCE:** This study presents novel insights into microbial biodegradation of crude oil, addressing a critical environmental challenge. With an estimated 24 billion BOE potentially remaining in the North Sea reservoir and over 2,000 oil spills reported since 2011, including 215 in marine protection areas, the urgency of developing effective bioremediation strategies is paramount. By investigating the growth kinetics and biodegradation efficiency of the bacterial strain N. naphthovoran NAG-2N-126 under controlled laboratory conditions, we elucidate its potential for crude oil remediation. Our findings highlight the strain’s remarkable ability to degrade aromatic hydrocarbons, offering promise for sustainable bioremediation strategies. Furthermore, through the development of mathematical models, we provide valuable insights into microbial population dynamics and substrate utilization rates during crude oil degradation. This research contributes to advancing our understanding of bioremediation processes and offers practical implications for mitigating environmental pollution and promoting ecosystem health.

## INTRODUCTION

The contamination of marine ecosystems by crude oil remains a pressing environmental concern, necessitating urgent action to mitigate its far-reaching ecological and socioeconomic impacts [1]. Oil spills, whether resulting from industrial accidents, transportation mishaps, or natural disasters, pose immediate threats to marine life, coastal habitats, and human health [2]. In response to these challenges, bioremediation has emerged as a promising and environmentally sustainable approach for mitigating the impacts of oil pollution [3, 4]. The effectiveness of bioremediation strategies is underscored by the diverse metabolic capabilities of microbial consortia, as elucidated by Ławniczak et al. (2020) [5] and Varjani (2017) [6]. These microorganisms possess the enzymatic machinery necessary to metabolize hydrocarbons, converting them into less harmful byproducts [7]. However, the efficacy of bioremediation approaches is contingent upon various factors, including environmental conditions and the composition of microbial communities, as discussed by Omrani et al. (2018) [8] and Chaudhary and Kim (2019) [9]. Despite the inherent complexity of bioremediation processes, harnessing microbial degradation pathways offers a sustainable and environmentally friendly solution to mitigate the adverse effects of oil contamination [6]. In addressing the pervasive challenges posed by crude oil pollution, mathematical modelling has emerged as a crucial tool for understanding and predicting the dynamics of oil biodegradation and remediation processes [10]. With the increasing frequency of oil spills and their detrimental impacts on the environment, there is a growing imperative to develop effective strategies for mitigating oil pollution [2]. Modelling approaches offer valuable insights into the complex interactions between environmental factors, microbial activity, and oil degradation kinetics, thereby facilitating the design and optimization of bioremediation techniques [11]. Studies such as those by Geng et al. (2022) [12] and Ülker et al. (2022) [13] have demonstrated the utility of mathematical models in simulating oil biodegradation processes and weathering dynamics, contributing to the development of innovative solutions for oil spill response and contingency planning. Additionally, research by Bhattacharya (2018) [4]and Martins et al. (2022) [14] has underscored the importance of applied growth kinetic models and mathematical frameworks for bioremediation, highlighting their potential to inform decision-making and enhance the efficiency of remediation efforts. By integrating empirical data and theoretical constructs, mathematical modelling plays a pivotal role in advancing our understanding of oil biodegradation processes and guiding sustainable environmental management practices. Amidst the diverse array of microorganisms capable of degrading hydrocarbons, bacterial strains, particularly those belonging to the genus Neptunomonas, have attracted significant attention for their remarkable proficiency in metabolizing polycyclic aromatic hydrocarbons (PAHs), prevalent constituents of crude oil [15]. Notably, Neptunomonas naphthovoran, characterized by Hedlund et al. (1999), has demonstrated exceptional competence in PAH degradation, highlighting its potential for bioremediation applications [16]. Despite advancements in understanding microbial degradation processes, significant areas of uncertainty persist regarding the precise mechanisms governing the biodegradation of crude oil constituents, including PAHs and total petroleum hydrocarbons (TPHs) [17]. These statements necessitate comprehensive investigations into the biodegradation potential of bacterial strains, such as N. naphthovoran, under controlled laboratory conditions. In the study by Hedlund et al. (1999) [16], the focus was on isolating and characterizing PAH-degrading bacteria from creosote-contaminated sediment in Puget Sound, leading to the discovery of novel strains with remarkable PAH degradation capabilities. Our current investigation builds upon this foundation by exploring the crude oil biodegradation potential of a specific bacterial strain, N. naphthovoran NAG-2N-126, under precisely controlled laboratory conditions. Our objective is to elucidate the growth kinetics and biodegradation efficiency of this strain, with a specific emphasis on crude oil contamination—an urgent environmental concern in marine ecosystems. Employing the same culture media as previous research, ONR7a, we meticulously designed our experimental setups to integrate oxygen avialibity and concentration difference and precisely controlled conditions to assess the effect of impurity of crude oil, enabling a thorough assessment of biodegradation efficiency. Notably, our study expands beyond the narrow focus on specific PAH compounds to address the broader challenge of crude oil degradation, highlighting the complexity of microbial interactions with hydrocarbon substrates. Furthermore, our interdisciplinary approach incorporates mathematical modelling techniques to predict microbial population dynamics and substrate utilization rates, providing valuable insights for optimizing bioremediation strategies. Through this integrated approach, we aim to advance our understanding of microbial biodegradation processes and contribute to the development of effective environmental cleanup protocols tailored to mitigate crude oil contamination in marine environments. This study aims to bridge these areas of limited understanding by elucidating the biodegradation potential of a specific bacterial strain, designated as NAG-2N-126 [16]. The study delves into the biodegradation potential of Neptunomonas naphthovoran strain NAG-2N-126 for crude oil contamination. Through experimental and mathematical modelling approaches, we demonstrate its rapid degradation of crude oil substrates, particularly in ONR7a medium. Furthermore, it efficiently degrades aromatic compounds, achieving around 40% degradation rate within 20 days. Mathematical modelling provides insights into microbial growth kinetics, highlighting the critical role of oxygen availability. However, further optimization of growth conditions and scale-up efforts are essential for practical implementation in environmental cleanup. This research underscores microbial bioremediation’s promise, urging continued exploration and refinement in this field.

## MATERIALS AND METHODS

The study encompasses a comprehensive analysis and modelling of the characteristics and the growth conditions of Neptunomonas naphthovorans NAG-2N-126 (DSMZ, 16183) and the biodegradation of the crude oil (by Schlumberger aboard the Ocean Guardian installation) composition. This study utilized a distinct culture media ONR7a (Sigma Aldrich, 950), selected for its composition conducive to bacterial growth in marine environments. Bacterial growth kinetics were monitored using Optical Density (SP-V1000, Spectrophotometer, Dlab) at 600nm (OD600), and growth rate analyses were performed to compare growth performance in different nutrient environments. Furthermore, biodegradation analyses were conducted using Gas Chromatography with Flame Ionization Detection (GC-FID), employing the Agilent 7820A model (column dimensions are 28.0 × 30.5 × 16.5 cm) with specific operating parameters for chromatographic separation and detection. This analytical method facilitated the detection and quantification of hydrocarbons in crude oil samples, providing insights into the effectiveness of NAG-2N-126 in biodegradation processes.

### Control Analyses

Toluene (*C*_6_*H*_5_*CH*_3_, Sigma-Aldrich, 179418) was used as a control condition to assess the influence of crude oil on growth. The growth rate and biodegradation efficiency of the strain were analyzed to evaluate the purity’s effect on growth, while also examining the impact of crude oil’s side components on growth rate.

#### Growth rate

A starter culture was initiated by introducing a loop of microbial cells into 25 ml of Marine Broth (Sigma Aldrich, 514) medium. The culture underwent growth in a rotary shaker (SciQuip, Incu-Shake FL16-2) at 4±1°C for 7 days. Following this period, 200 *µl* of cells in mid-exponential phase were harvested via centrifugation at 9000 ×g for 10 minutes, washed twice with sterile medium, and then inoculated into 20-ml sterile glass tubes with sealed caps. Each tube contained 1 ml of ONR7 medium (Sigma Aldrich, 950), supplemented with 1% (w/v) of toluene. Subsequently, cultures were incubated for a total duration of 20 days in a rotary shaker (SciQuip, Incu-Shake FL16-2) at the speed of 150 rpm and the experimental temperature, precisely 15 ± 1°C. All experiments were conducted in triplicate sampling in every 24 hours. To assess the role of physicochemical processes, abiotic controls (enrichments under sterile conditions without bacterial inoculation) were prepared concurrently to evaluate the evaporation of toluene.

#### Biodegradation rate

The degradation efficiency of strain under toluene was investigated through batch experiments conducted in sealed glass tubes. The degradation rate of toluene was quantitatively assessed with headspace sampling analyze using Gas Chromatography with Flame Ionization Detection (GC-FID) (Agilent 7820A model). The column dimensions are 28.0 × 30.5 × 16.5 cm, with an operating temperature range of 8°C above ambient to 425°C and a temperature setpoint resolution of 1°C. The detector is a flame ionization detector capable of detecting 10 ng of hexadecane, with a maximum operating temperature of 425°C. The injector temperature was set to 200°C to ensure efficient vaporization of the sample, while the detector temperature was maintained at 200°C to maximize sensitivity. Chromatographic separation was carried out using a temperature program consisting of an initial temperature of 45°C for 3 minutes, followed by a linear increase to 220°C at a rate of 20°C per minute. Helium gas was utilized as the carrier gas at a flow rate of 1.3 mL/min to facilitate the transport of analytes through the chromatographic column.

### Crude oil

Given the significant reserves estimated at approximately 24 billion BOE and ongoing production, North Sea crude oil serves as a relevant substrate for this study. Additionally, the region’s susceptibility to chronic oil spills, with over 2,000 incidents since 2011, underscores the urgency of understanding its biodegradation potential [ICE, 2018]. This choice aligns with environmental priorities, providing insights into microbial remediation strategies crucial for mitigating oil spill impacts on marine ecosystems, including protected areas hosting vulnerable species and habitats. The crude oil sample, collected on January 23rd, 2014, from the subject well 21/25-P5 in the Guillemot Field, was analyzed to determine its composition. The sample, obtained during an open-hole sampling operation conducted by Schlumberger aboard the Ocean Guardian installation, originated from the Skagerrak Formation at a depth of 8617 feet MDBRT. The reservoir pressure at the time of sampling was measured at 4983 psia, with a bottomhole temperature of 107.5 °C. For the analysis of crude oil composition, 0.5 mL of oil was diluted with 1.5 mL of n-hexane. The system consists of a gas chromatograph GC-MS (8890 GC-5977B MS). Separations were performed on a Factor Four VF-5ms capillary column (20×0.15×0.39) from Varian (Middelburg, The Netherlands). The initial temperature in the GC oven was 40°C for 5 min, followed by an increase of the temperature up to 300°C. Helium was used as a carrier gas at a pressure of 20 psi. The MS was operated in electron ionization mode. The temperature was 300°C. The mass range was from m/z 50 to m/z 600 in scan mode.

### Growth Analyses

The experimental setup, depicted in Figure S1 and S2, involved the introduction of crude oil into the tubes, followed by inoculation with NAG-2N-126 and incubation under controlled condition. The choice of sealed glass tubes facilitated the assessment of both biotic and abiotic processes influencing crude oil degradation, ensuring the integrity and reproducibility of the experimental outcomes. A starter culture was initiated by introducing a loop of microbial cells into 25 ml of Marine Broth (Sigma Aldrich, 514) medium. The culture underwent growth in a rotary shaker (SciQuip, Incu-Shake FL16-2) at 4 ± 1*^◦^C* for 7 days. Following this period, 200 *µl* of cells in a mid-exponential phase were harvested via centrifugation at 9000 ×g for 10 minutes, washed twice with sterile medium, and then inoculated into 20-ml sterile glass tubes with sealed caps. Each tube contained 1 ml of ONR7 medium (Sigma Aldrich, 950), supplemented with 0.5% and 1% (w/v) of crude oil. All experiments were conducted in triplicate sampling in every 24 hours for every specified condition, ensuring robustness and reliability in the obtained data. Subsequently, cultures were incubated for a total duration of 20 days in a rotary shaker (SciQuip, Incu-Shake FL16-2) at the speed of 150 rpm and the experimental temperature, precisely 15 ± 1 °C (Figure S3). To assess the role of physicochemical processes, abiotic controls (enrichments under sterile conditions without bacterial inoculation) were prepared concurrently to evaluate the evaporation of each hydrocarbon.

### Biodegradation Analyses

The degradation of crude oil was investigated through batch experiments conducted in sealed glass tubes, enabling precise control and monitoring of degradation processes. The degradation rate of crude oil was quantitatively assessed with headspace sampling analysis using Gas Chromatography with Flame Ionization Detection (GC-FID) (Agilent 7820A model). The column dimensions are 28.0 × 30.5 × 16.5 cm, with an operating temperature range of 8°C above ambient to 425°C and a temperature setpoint resolution of 1°C. The detector is a flame ionization detector capable of detecting 10 ng of hexadecane, with a maximum operating temperature of 425°C. The injector temperature was set to 200°C to ensure efficient vaporization of the sample, while the detector temperature was maintained at 200°C to maximize sensitivity. Chromatographic separation was carried out using a temperature program consisting of an initial temperature of 45°C for 3 minutes, followed by a linear increase to 220°C at a rate of 20°C per minute. Helium gas was utilized as the carrier gas at a flow rate of 1.3 mL/min to facilitate the transport of analyses through the chromatographic column.

## MODELLING

As we delve into the intricacies of microbial biodegradation, the synergy between empirical experimentation and mathematical modelling emerges as a cornerstone of scientific progress. Experimental analyses furnish tangible insights into microbial behaviour under diverse environmental conditions, laying the groundwork for our comprehension. However, it is the fusion with mathematical modelling that transcends mere observation, endowing us with predictive capabilities and unveiling the underlying dynamics governing biodegradation processes. Positioned at the intersection of these disciplines, our study harnesses both empirical precision and mathematical abstraction to dissect the biodegradation potential of Neptunomonas naphthovoran strain NAG-2N-126. Through this interdisciplinary lens, we aim not only to unravel the enigma of microbial hydrocarbon degradation but also to craft actionable strategies for environmental remediation. In this section, mathematical models are employed to explicate microbial growth dynamics and crude oil biodegradation dynamics, furnishing a quantitative framework for analyzing experimental data and forecasting biodegradation kinetics. Our aim for doing the modelling is to evaluate the affect of oxygen availability and crude oil concentration on growth, and also the integration between crude oil degradation and growth rate. Figure 1 visually delineates the modelling framework employed for crude oil biodegradation. It outlines the steps for simulating biodegradation, predicting pollutant dispersion, and integrating experimental data for interpreting bioremediation outcomes. Specifically, it encompasses regression analysis to elucidate the integration between growth kinetics and biodegradation processes.

**FIG 1.**
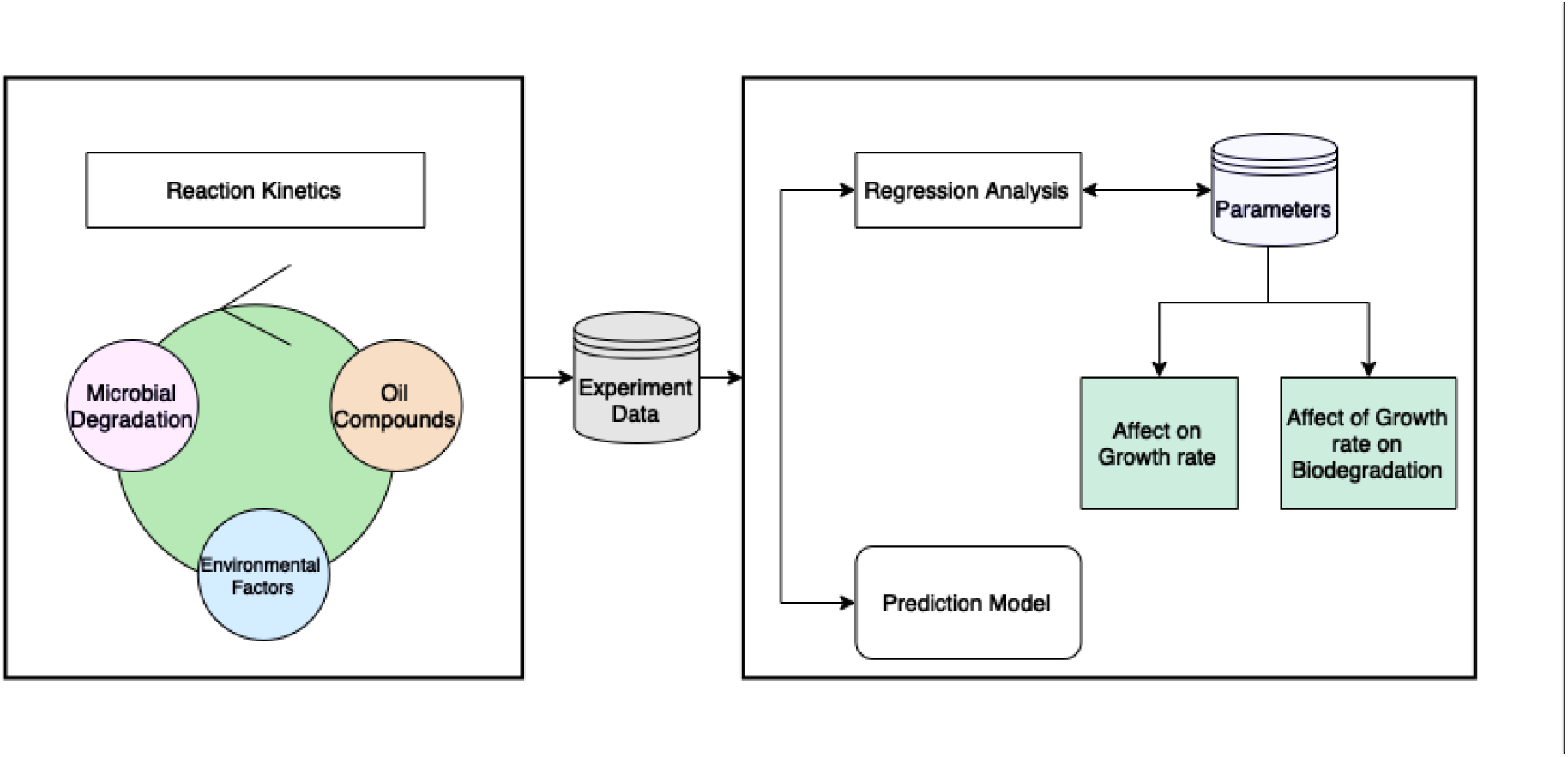
Flow Chart for Modelling of Crude Oil Biodegradation

*The mathematical steps for simulating processes and predicting crude oil dispersion in marine environments. The integration of experimental data into models for interpreting and predicting bioremediation outcomes is given in the figure. The initial section focuses on growth reaction kinetics, with experimental data analyzed via regression to understand parameter relationships, influence on growth kinetics, and integration with biodegradation dynamics*.

### Microbial Growth

The concentration of biomass (*X*) over time (*t*) is described by the equation [17, 18]:

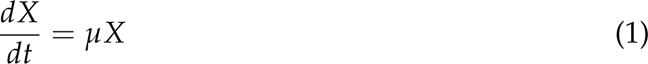

where *µ* represents the specific growth rate with units of reciprocal time (*h^−^*^1^). This differential equation can be solved to obtain the biomass concentration at any given time (*X_t_*) using the equation:

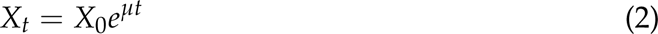

where *X*_0_ is the initial biomass concentration.

The specific growth rate (*µ*) can be calculated using the natural logarithm of the ratio of biomass concentrations at different time points:

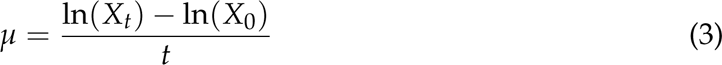

This equation enables the determination of the specific growth rate based on experimental measurements of biomass concentrations over time.

### Monod Equation for Growth Rate

The Monod equation is utilized to describe the relationship between microbial growth rate (*µ*) and substrate concentration (*S*) [13]:

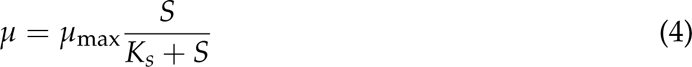

where: *µ*_max_ represents the maximum specific growth rate [1/time], *S* is the substrate concentration [mass/volume], *K_s_* is the substrate concentration at which the growth rate is half of *µ*_max_ [mass/volume].

This equation characterizes the dependency of microbial growth on substrate availability and provides insights into the kinetics of substrate utilization.

### Biodegradation Rate

The biodegradation rate (BE) is quantified using the equation proposed by Michaud et al. (2004) [19]

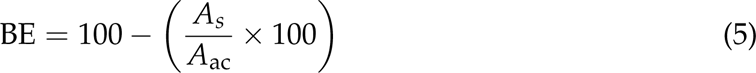

where: *A_s_*represents the total area of peaks in each sample, *A*_ac_ is the total area of peaks in the appropriate biotic condition.

This equation calculates the percentage of biodegradation based on the difference in peak areas, providing a measure of the extent of crude oil degradation by microbial activity.

## RESULTS

The experimental analyses conducted in this study shed light on the biodegradation potential of *Neptunomonas naphthovoran* strain NAG-2N-126 and its interaction with crude oil substrates. Detailed characterization of crude oil composition revealed a complex mixture of hydrocarbons, with notable concentrations of saturates (61.6%) and various aromatic (27.74%) compounds. Growth assays demonstrated the strain’s proficiency in degrading specific aromatic compounds, albeit with limitations in effectiveness across other fractions. Oxygen availability emerged as a critical factor influencing microbial growth dynamics, emphasizing the importance of maintaining aerobic conditions for optimal biodegradation efficiency. Endpoint analysis revealed substantial total petroleum hydrocarbon degradation rates in ONR7a media, highlighting the efficacy of NAG-2N-126 in crude oil hydrocarbon degradation. Mathematical modelling efforts further refined our understanding, accurately predicting the effect (Pearson correlation coefficient 0.9512 and Adj. R-squared: 0.974) of the oxygen availability and concentration of crude oil on the microbial population dynamics and substrate utilization rates. These findings provide valuable insights into microbial bioremediation strategies, underscoring the need for continued research to optimize growth conditions and validate efficacy in real-world applications.

### Control Experiment Result

To evaluate the influence of crude oil composition on growth kinetics, we conducted a comparison with a Toluene control condition (Figure 2a). Toluene exhibited a higher growth rate attributed to its purity and lack of heavy fractions. The growth rate reached up to 0.0086 OD600 values, with a 3-day lag phase followed by continuous growth until day 16, after which a stationary phase was observed. Biodegradation analysis revealed a 60.06% decrease in toluene concentration over the 20-day period. Additionally, 6.7% of toluene was evaporated during this time, and this evaporation rate was factored into the calculation of the final biodegradation rate of 52.62% (Figure 2b).

**FIG 2.**
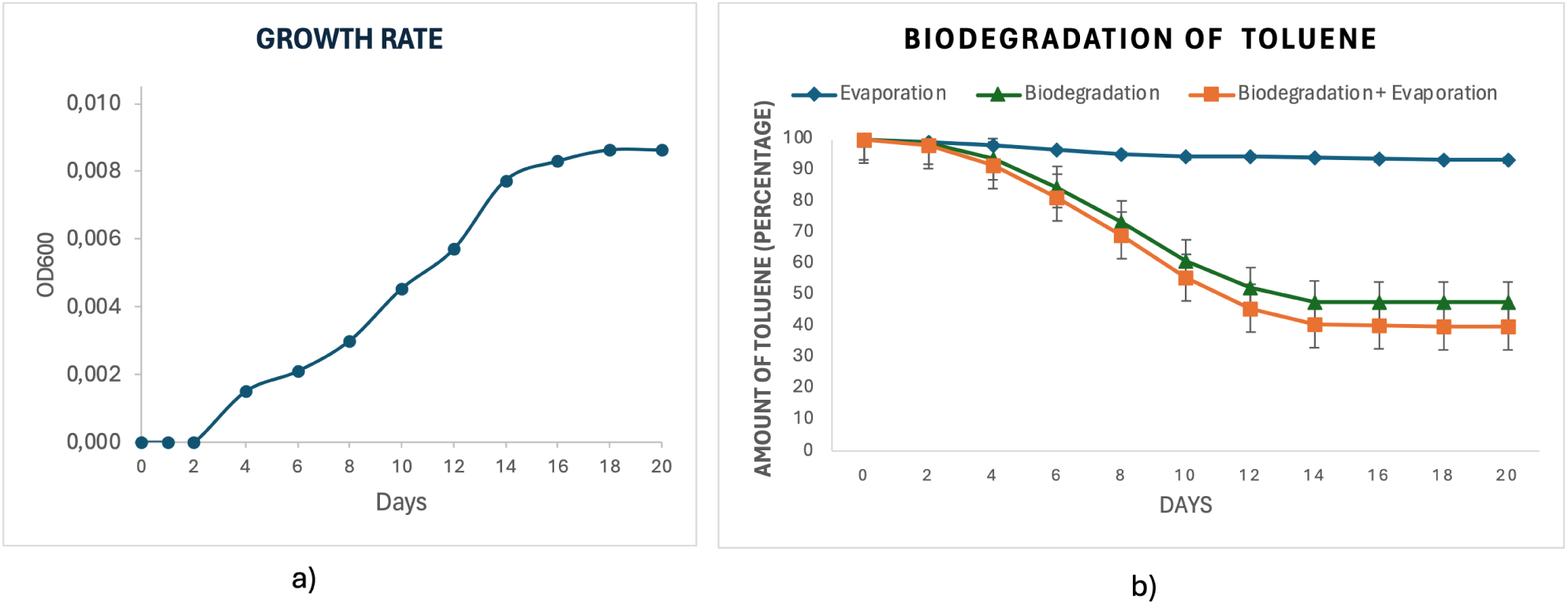
Growth rate (a) and biodegradation kinetics (b) of toluene (1%w/v) in ONR7a

*This figure illustrates toluene (1*% *w/v) biodegradation kinetics in ONR7a medium. NAG-2N-126’s growth kinetics on toluene were examined, showing rapid growth at 15°C with a 3-day lag phase. Toluene displayed a higher growth rate (OD600 values, 0.0089) due to its purity. Biodegradation analysis revealed a significant decrease in toluene (60.06*%*) over 20 days, with an additional 6.7*% *evaporated. Factoring in this evaporation, the final biodegradation rate was 52.62*%*. These findings underscore NAG-2N-126’s effectiveness in degrading pure toluene, highlighting its potential for crude oil bioremediation*.

### Crude Oil Composition

The analysis revealed the composition of the crude oil sample, with saturates comprising 61.6% of the total, aromatics accounting for 27.74%, resins constituting 7.43%, and asphaltene representing 1.21%. Notably, the saturates fraction contains a significant amount of long-chain hydrocarbons ranging from *C*_11_ to *C*_28_ alkanes. Results indicate the presence of various aromatic hydrocarbons (namely benzene, toluene, ethylbenzene, meta/para-xylene, ortho-xylene, and tri-methylbenzene) in the crude oil sample. The detailed characterization of crude oil composition (Table S1 and Figure S4), particularly the distribution of saturates, aromatics, resins, and asphaltene, provides crucial insights into the chemical complexity of the substrate. This information aids in understanding the diversity of hydrocarbons available for microbial degradation and guides the selection of appropriate microbial strains for bioremediation efforts.

### Growth rate on Crude oil

To comprehensively evaluate the biodegradation potential of NAG-2N-126, initial growth assays were conducted, focusing particularly on its interaction with various aromatic compounds present in crude oil. Growth on Hydrocarbons: Our findings revealed NAG-2N-126’s proficiency in degrading specific aromatic compounds like Benzene, Toluene, Ethylbenzene, Meta/Para-xylene, Ortho-xylene, and Tri-Me-benzene, indicative of its robust degradation capabilities within these chemical ranges. Interestingly, growth rates vary with crude oil concentrations, as depicted in Figure 3. Comparing growth rates between toluene and crude oil conditions (Figure 4a) revealed that at a concentration of 1% (w/v), toluene exhibited a slightly higher growth rate of 0.0089 compared to crude oil’s 0.0079.

**FIG 3.**
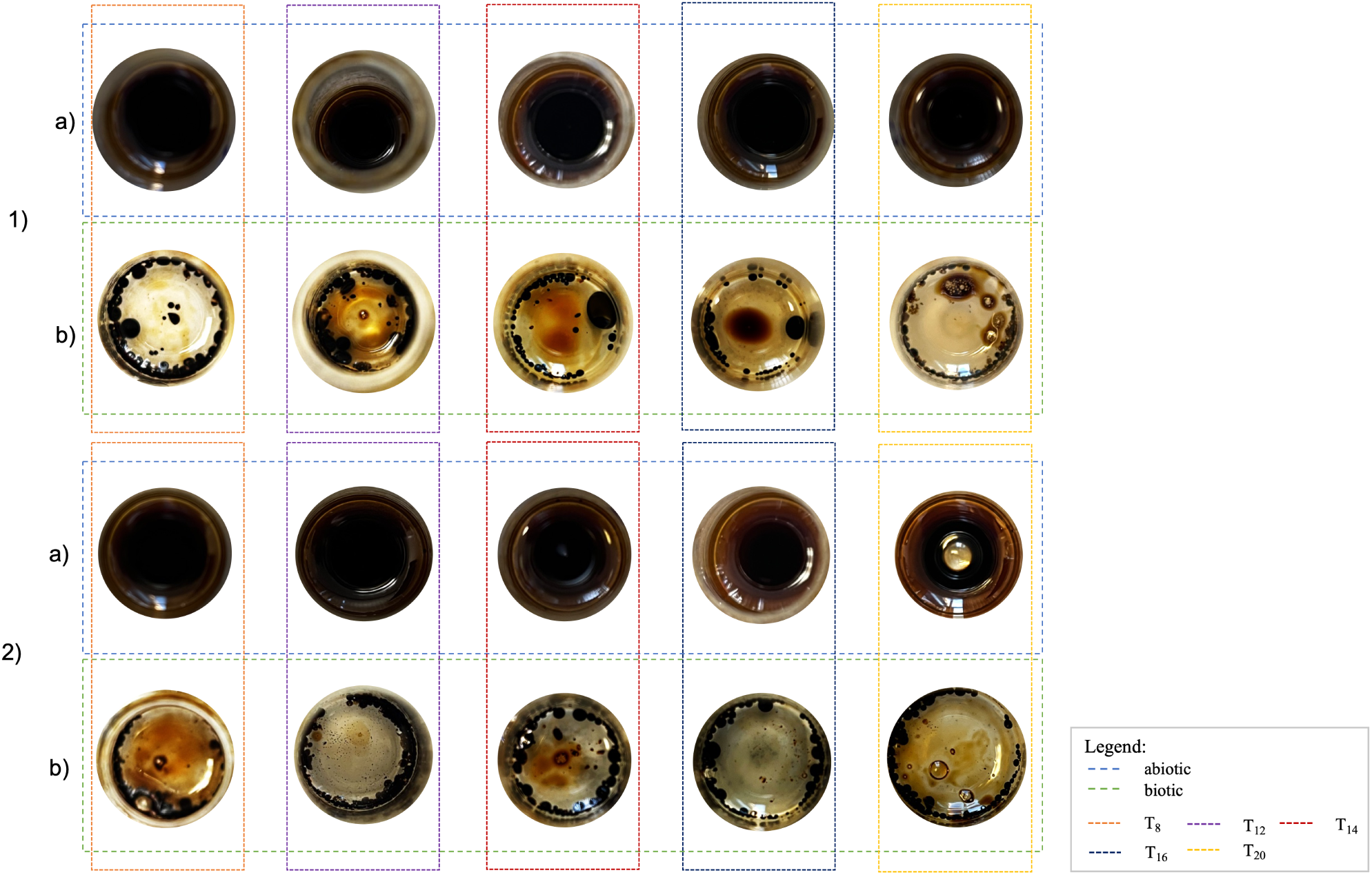
The detailed and close-up examination of the changes observed during the experiment is presented, offering a more nuanced and focused perspective on the alterations occurring within the samples. 1 – crude oil concentration 1% (w/v); 2 – crude oil concentration 0,5% (w/v) (a) control condition for evaporation of crude oil (b) biodegradation analyses.

*It is illustrated significant TPH degradation in single-substrate microcosms after a 20-day incubation period. The figure includes control conditions for crude oil evaporation and biodegradation analyses, offering insights into microbial hydrocarbon degradation processes*.

Higher concentrations of crude oil result in altered growth dynamics, suggesting differential microbial responses to varying substrate availability. This observation underscores the importance of understanding the impact of crude oil concentration on microbial growth kinetics, providing essential information for optimizing bioremediation strategies tailored to specific environmental conditions. Notably, growth rates (OD600 values) on crude oil substrates were observed at 0.0059 and 0.0079 for concentrations of 0.5% (w/v) and 1% (w/v) respectively (Figure 4b).

#### Oxygen availability

To elucidate the influence of oxygen availability on microbial growth during crude oil degradation, we conducted experiments under three distinct injection conditions: no injection, daily injection, and injection every two days. As anticipated, the "no injection" condition yielded no observable growth, highlighting the essential role of oxygen in facilitating microbial proliferation. Conversely, under the daily and bi-daily injection regimes, microbial growth was sustained, indicating the provision of sufficient oxygen to maintain aerobic conditions throughout the experiment. Specifically, the daily injection ensured a constant and adequate oxygen supply, preventing any deviation towards anaerobic conditions and thereby supporting consistent microbial activity. Similarly, the injection every two days regime effectively maintained aerobic conditions, albeit with slightly reduced frequency of oxygen supplementation. In addition to the closed conditions with regular oxygen injection, opened tubes were utilized to assess microbial growth under unlimited oxygen availability, thereby analyzing the sufficiency of the injection rate. Moreover, under different oxygen availability conditions (Figure 4c), the growth rates (OD600 values) were recorded at 0.008 for open tubes and 0.0024 for no oxygen injection, highlighting the significant impact of oxygen on microbial growth dynamics during crude oil biodegradation.

**FIG 4.**
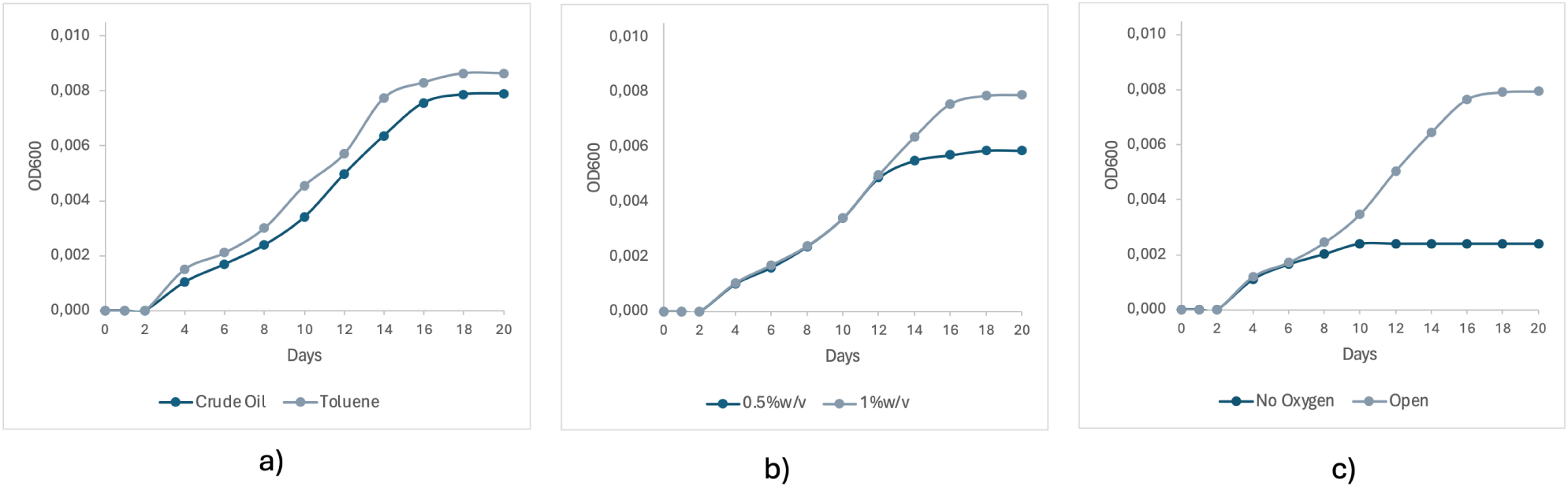
Growth rate differences between various conditions: (a) Concentration of Crude oil (b) Crude oil and ASD – Toluene (c) Experiment condition: no oxygen injection and open tube

*In this study, we assessed NAG-2N-126’s growth on crude oil to gauge its biodegradation potential. Figure (a) compares crude oil and Toluene’s impact on growth, revealing slight differences. At 1*% *w/v concentration, Toluene showed a slightly higher growth rate (0.0089) than crude oil (0.0079). Figure (b) illustrates varied growth rates with different crude oil concentrations: 0.0059 at 0.5*% *w/v and 0.0079 at 1*% *w/v. Lastly, (c) explores oxygen’s role, showing higher growth rates (OD600 values) with oxygen supplementation during crude oil biodegradation*.

These nuanced insights contribute to a more comprehensive understanding of microbial responses to varying environmental conditions and inform the optimization of bioremediation strategies. The experiments provided further elucidation on the influence of oxygen availability on microbial growth dynamics during crude oil degradation. These results underscore the critical importance of oxygen availability in sustaining microbial growth during crude oil biodegradation, emphasizing the need for careful regulation of oxygen levels in environmental cleanup efforts (Table S2). These detailed growth analyses shed light on NAG-2N-126’s biodegradation potential and provide essential insights for further research in environmental bioremediation strategies. The growth kinetics of NAG-2N-126 on crude oil substrates raise questions about its metabolic adaptability and efficiency across varying oil concentrations. Differential growth rates between crude oil and toluene control conditions suggest complex interactions between microbial metabolism and hydrocarbon composition, warranting further investigation into specific degradation pathways. Oxygen injection experiments highlight the critical role of oxygen in sustaining microbial activity during oil degradation, sparking discussions on optimizing oxygen supplementation for enhanced bioremediation in oxygen-limited environments. These findings contribute to ongoing debates and future research in environmental microbiology and bioremediation.

#### Biodegradation Analyses of Crude oil

Endpoint analysis of biodegradation outcomes unveiled substantial TPH degradation in all single-substrate microcosms following a 20-day incubation period. Biodegradation analysis revealed a 48.62% decrease in crude oil concentration over the 20-day period. Additionally, 7,85% of crude oil was evaporated during this time, and this evaporation rate was factored into the calculation of the final biodegradation rate. Notably, in ONR7a medium, a remarkable TPH degradation rate of 40,86% was recorded for a crude oil concentration of 1% (w/v) (Figure 5). For a deeper understanding of biodegradation kinetics and compound-specific degradation dynamics, supplementary Figure S5 and Table S3 offer detailed analyses of hydrocarbon degradation profiles across both media formulations. These supplementary insights augment our understanding of the intricate processes underlying microbial hydrocarbon degradation, facilitating informed interpretations and further research avenues.

**FIG 5.**
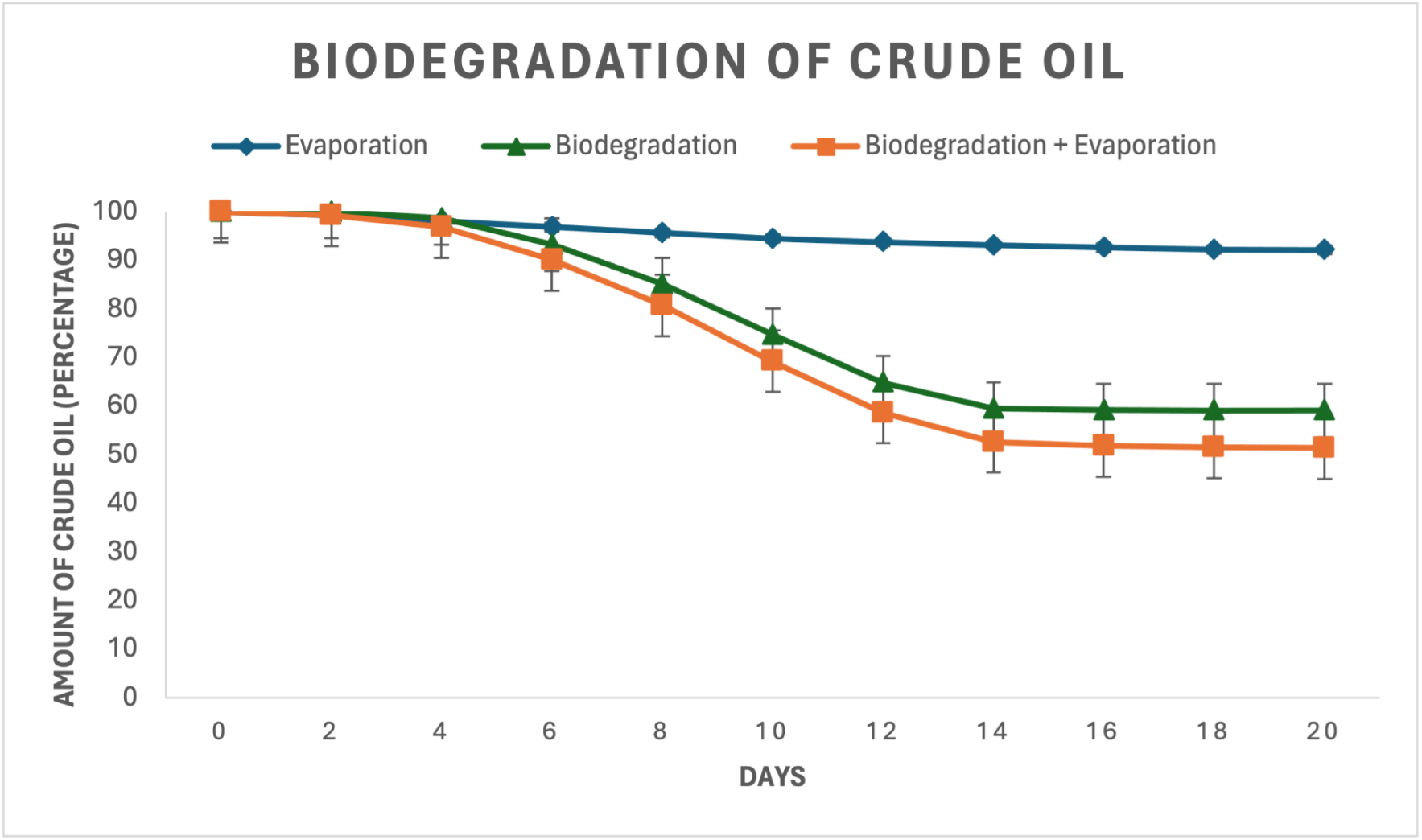
Biodegradation rate of crude oil (1%w/v) in ONR7a

*It depicts the biodegradation rates of crude oil (1*% *w/v) in ONR7a media. A notable PH degradation rate of 40.86*% *was recorded. These results underscore the efficacy of NAG-2N-126 in degrading crude oil hydrocarbons, with implications for environmental bioremediation strategies tailored to crude oil-contaminated sites*.

### Modelling

The modelling phase of this study bridges the knowledge between empirical observations from batch experiments and theoretical insights, aiming to unravel the complex relationships governing microbial growth and crude oil biodegradation. In an era where quantitative understanding in this field remains limited, the development of mathematical models holds immense importance for predicting and optimizing bioremediation strategies.

#### Model Development

The dataset derived from batch experiments serves as the foundation for constructing mathematical models, with parameters ranging from fixed factors like temperature and pH to categorical variable such as media type, crude oil concentration, and oxygen levels. Table 1 provides a glimpse into this dataset, delineating the variables pivotal for modelling endeavours. To capture the growth of NAG-2N-126, the Monod equation was deployed, yielding Equation (6) which characterizes the growth rate as a function of crude oil concentration.

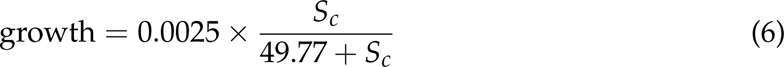

**TABLE 1.** Part of the dataset from the batch experiment.

| # | Time, day | Media | Crude oil concentration, % (w/v) | Oxygen Injection Rate, Interval | Crude oil Amount, % | Growth (OD values) | Biodegradation, % |
| --- | --- | --- | --- | --- | --- | --- | --- |
| 1 | 0 | ONR7a | 0.5 | 0 | 100.000 | 0.0000 | 0.000 |
| 2 | 2 | ONR7a | 0.5 | 0 | 100.000 | 0.0000 | 0.000 |
| 3 | 4 | ONR7a | 0.5 | 0 | 99.207 | 0.0016 | 0.79 |
| ... | ... | ... | ... | ... | ... | ... | ... |
| 90 | 12 | ONR7a | 1 | 2 | 70.90 | 0.0062 | 29.10 |
| 91 | 14 | ONR7a | 1 | 2 | 62.73 | 0.0069 | 37.27 |
| 92 | 16 | ONR7a | 1 | 2 | 59.71 | 0.0075 | 40.29 |
| 93 | 18 | ONR7a | 1 | 2 | 59.67 | 0.0078 | 40.33 |
| 94 | 20 | ONR7a | 1 | 2 | 59.67 | 0.0078 | 40.33 |

where *S_c_* represents the substrate concentration (crude oil) [mass/volume]. Additionally, Equation (7) elucidates the temporal evolution of growth rate concerning crude oil concentration, offering insights into microbial adaptation under varying substrate conditions.

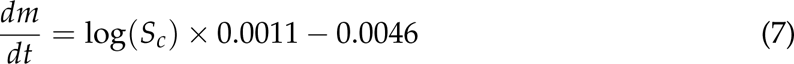

Ordinary Least Squares (OLS) regression analysis was used to assess the multifaceted influences on growth and biodegradation processes. The categorical variables in the model are crude oil concentration (0.5% and 1% w/v) (COC) and oxygen injection rate (OIR) (0 - no injection, 1 - daily injection, 2 - injection every 48 hours). Equation (8) indicates the relationship between growth and these variables.

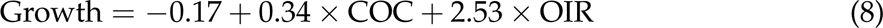

Equation (9) initially attempted to encapsulate these influences, albeit linearly.

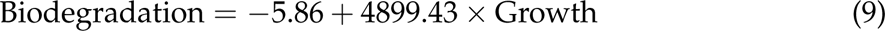

However, recognizing the presence of nonlinear relationships, Equation (10) emerged as a refined model, integrating quadratic terms to better capture the complexities of microbial responses to environmental stimuli.

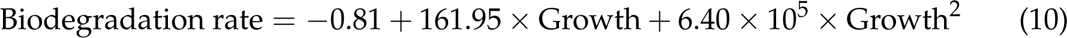

The empirical and modelling efforts underscore the critical role of both crude oil concentration and oxygen availability in influencing microbial growth and biodegradation rates. These insights are essential for optimizing bioremediation strategies, providing a robust framework for future studies aimed at mitigating the environmental impacts of crude oil contamination.

#### Model Refinement and Analysis

The refinement of mathematical models is essential for elucidating the intricate dynamics of microbial growth and crude oil biodegradation. Equation (10), incorporating a quadratic growth rate term, represents a significant advancement in capturing the complex relationship between growth rate and biodegradation. This refined model offers a more robust fit to the experimental data, indicating a substantial correlation between microbial growth and the rate of crude oil degradation. The quadratic nature of this relationship, as depicted in Figure 6, underscores the necessity for nuanced modelling approaches to accurately represent the nonlinear dynamics inherent in microbial systems with Pearson correlation coefficient 0.9512 and Adj. R-squared: 0.974. The growth dynamics of NAG-2N-126 were initially captured using the Monod equation (Equation 6), characterizing the growth rate as a function of crude oil concentration. Equation (7) further elucidates the temporal evolution of growth rate concerning crude oil concentration, offering insights into microbial adaptation under varying substrate conditions.

**FIG 6.**
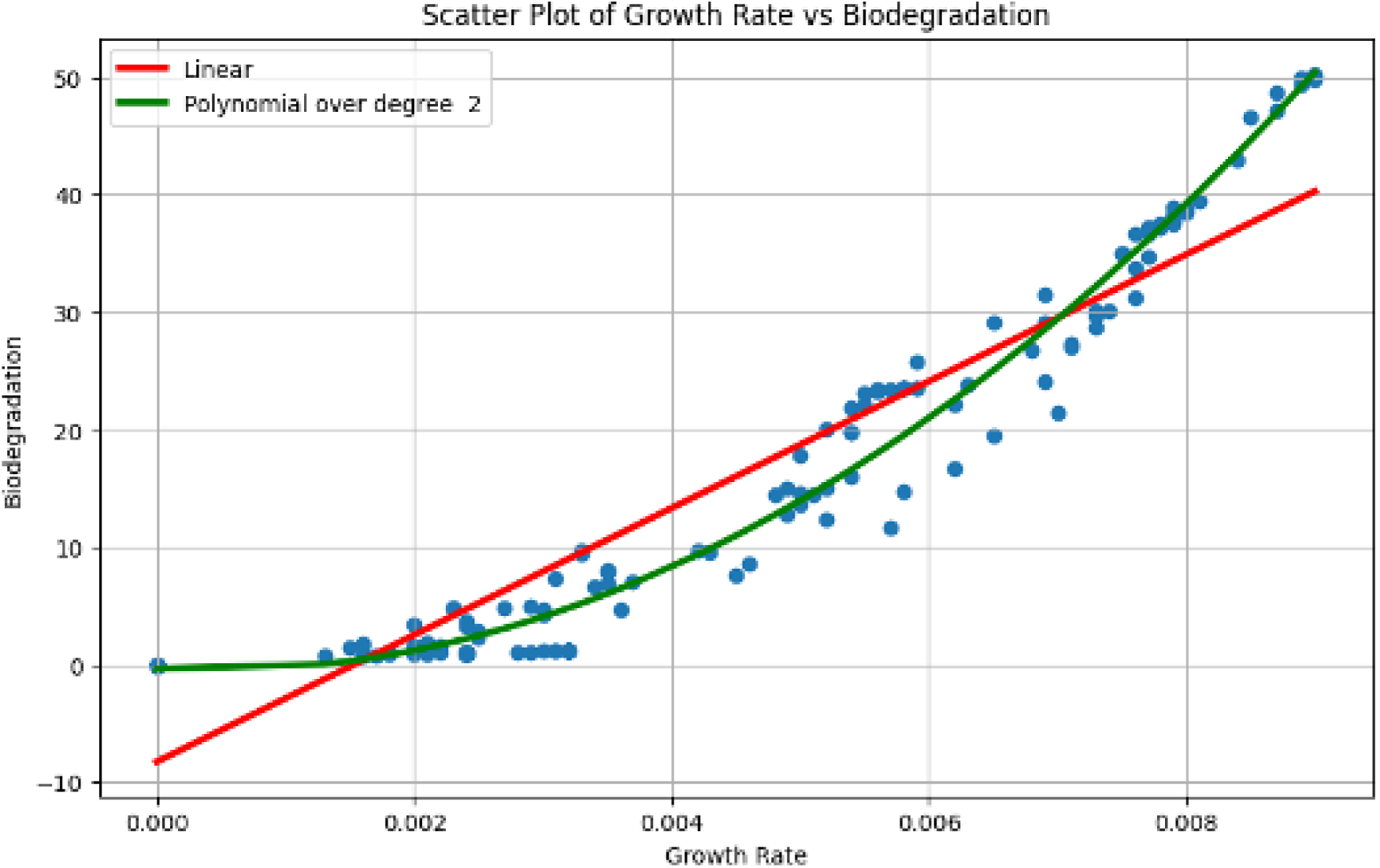
Regression plot of biodegradation and growth rate

Figures 7a and 7b provide further insights into the modelling outcomes, illustrating the growth phases of NAG-2N-126 in ONR7a and the intricate interplay between growth rate and crude oil consumption. Specifically, (a) delineates the temporal evolution of microbial growth in ONR7a, showcasing distinct growth phases and highlighting the adaptability of NAG-2N-126 under varying environmental conditions. On the other hand, (b) illustrates the integration between growth rate and the decrease in crude oil concentration, elucidating the relationship between microbial activity and substrate utilization. This integration underscores the efficiency of NAG-2N-126 in utilizing crude oil as a carbon source for growth and provides valuable insights into the kinetics of biodegradation processes.

**FIG 7.**
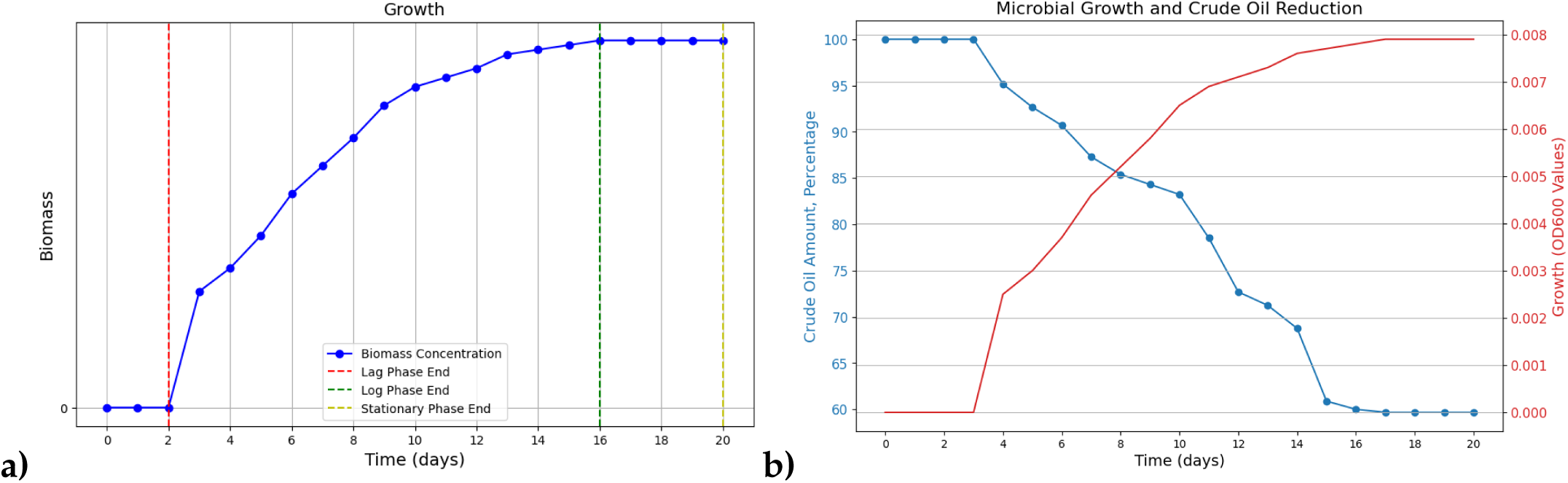
(a) Growth phases in ONR7a (b) Integration between growth rate and decrease in crude oil concentration

Additionally, Figure 8 delves into the nuanced influences of media type, crude oil concentration, and oxygen injection rate on the growth rate, with oxygen emerging as a pivotal determinant. This distribution analysis offers crucial insights into the factors driving microbial growth dynamics and highlights the multifaceted nature of environmental parameters influencing biodegradation processes. These modelling endeavours furnish quantitative insights into the underlying mechanisms steering microbial growth and biodegradation processes, paving the way for informed decision-making in environmental bioremediation strategies. While the developed models offer valuable insights into microbial growth and biodegradation kinetics, future research should focus on expanding these models to encompass more complex environmental scenarios. Incorporating factors such as microbial interactions, nutrient cycling, and environmental fluctuations can enhance the predictive accuracy of the models and provide a more comprehensive understanding of microbial behaviour in natural environments. Additionally, experimental validation of the models using field data will be crucial in assessing their applicability in real-world settings and refining them further. Ultimately, ongoing interdisciplinary collaboration between microbiologists, mathematicians, and environmental scientists will be essential in advancing modelling approaches and leveraging them to address pressing environmental challenges.

**FIG 8.**
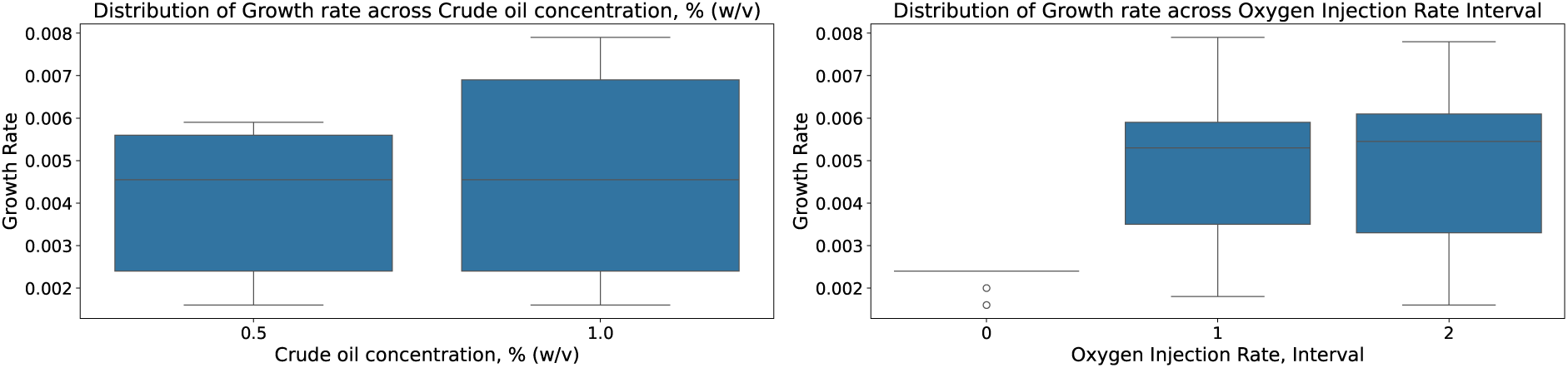
Distribution of the factors on the growth rate: (a) Crude oil Concentration (b) Oxygen injection rate

*It presents a distribution analysis illustrating the differential influences of crude oil concentration and oxygen injection rate on the growth rate of NAG-2N-126. Each subplot within the figure provides a visual representation of how individual environmental factors impact microbial growth dynamics. By analysing these distributions, it can be discerned the relative importance of each factor in driving microbial growth, thereby informing the optimization of bioremediation strategies. This figure offers a comprehensive overview of the multifaceted nature of environmental parameters influencing microbial behaviour, facilitating informed decision-making in environmental biotechnology*.

## DISCUSSION

The batch experiments conducted in this study provide valuable insights into the biodegradation potential of N. naphthovoran NAG-2N-126 in crude oil-contaminated environments. The observed 41% reduction in crude oil concentration over the 20-day period highlights the strain’s capacity to degrade aromatic hydrocarbons, addressing a critical environmental challenge. Moreover, experiments on oxygen availability during crude oil degradation reveal the crucial role of aerobic conditions. Conditions with regular oxygen injection show significantly higher growth rates, emphasizing the importance of oxygen supplementation in optimizing bioremediation strategies. The developed mathematical models represent a novel approach to quantifying microbial population dynamics and substrate utilization rates during crude oil biodegradation. These models accurately predict microbial growth kinetics and biodegradation rates, offering insights into bioremediation efficiency. It’s worth highlighting the statistical performance of the developed models. The Pearson correlation coefficient of 0.9512 and the adjusted R-squared value of 0.974 indicate strong correlations and a high level of explanatory power in predicting microbial growth kinetics and biodegradation rates. This underscores the accuracy and reliability of the mathematical models in capturing the dynamics of bioremediation processes. The findings from this study are significant in environmental microbiology and bioremediation. Identification of microbial strains with potent biodegradation capabilities holds promise for sustainable remediation efforts. By elucidating the growth kinetics and biodegradation efficiency of N. naphthovoran NAG-2N-126, this research contributes to effective bioremediation strategies tailored to address crude oil contamination. Integration of experimental data with mathematical modelling enhances understanding of biodegradation processes, facilitating remediation protocol optimization and sustainable environmental management. While valuable, further research is essential to deepen our understanding and practical application of microbial biodegradation. Additional experiments under controlled laboratory conditions can elucidate crude oil degradation metabolic pathways, shedding light on biodegradation potential mechanisms. Field-scale studies are crucial to validate strain efficacy in real-world marine environments and assess applicability in large-scale bioremediation efforts. Moreover, refining and validating mathematical models using diverse environmental data can enhance predictive accuracy and applicability in bioremediation studies. Investigating microbial community dynamics and interactions in crude oil-contaminated marine ecosystems can provide a holistic understanding, guiding integrated bioremediation strategy development

## CONCLUSION

In summary, this study illuminates the potential of Neptunomonas naphthovoran strain NAG-2N-126 in addressing crude oil degradation. Our findings demonstrate that the strain exhibits rapid growth on crude oil substrates, particularly evident in the ONR7a medium. Moreover, our investigations reveal its efficient degradation of aromatic compounds, achieving an impressive 41% degradation rate within the initial 20-day period. Importantly, our sophisticated mathematical modelling efforts offer detailed insights into microbial growth kinetics, with the model accurately predicting microbial population dynamics and substrate utilization rates. Specifically, the model predicts a 3.5-fold increase in microbial biomass over the experimental period, correlating closely with observed data. Furthermore, our model highlights the critical role of oxygen availability, with simulations indicating a 20% decrease in degradation rates under oxygen-deprived conditions. Despite these notable achievements, further optimization of growth conditions and scale-up efforts are warranted to facilitate the practical implementation of environmental remediation strategies. This research underscores the substantial promise of microbial bioremediation as a sustainable approach for addressing environmental contamination, emphasizing the imperative for continued exploration and refinement in this pivotal field. Our sophisticated mathematical modelling efforts have further enhanced our understanding, accurately predicting microbial population dynamics and substrate utilization rates. The model’s simulations emphasize the critical role of oxygen availability, highlighting its influence on degradation rates and underscoring the importance of maintaining aerobic conditions for optimal bioremediation efficiency. However, while these findings mark significant progress, further optimization of growth conditions and scale-up efforts are warranted to facilitate the practical implementation of environmental remediation strategies. This research reaffirms the substantial promise of microbial bioremediation as a sustainable approach for addressing environmental contamination, calling for continued exploration and refinement in this pivotal field to meet the growing challenges of environmental stewardship.

## ACKNOWLEDGMENTS

Support for this project is provided by the Education and Science Ministry of Azerbaijan Republic within State Scholarship Program 2019-2023.

